# Drought adaptability of different subspecies of tetraploid wheat (*Triticum turgidum*) under contrasting moisture conditions: association with solvent retention capacity and quality-related traits

**DOI:** 10.1101/2022.09.19.508520

**Authors:** F. Saeidnia, F. Shoormij, A. Mirlohi, E. Soleimani Kartalaei, M. Mohammadi, M. R. Sabzalian

## Abstract

Few prior efforts have been made to investigate the genetic potential of different subspecies of *Triticum turgidum* for drought tolerance and their quality-related traits compared with common wheat (*Triticum aestivum*) and to identify the association among agronomic, micronutrients, and quality-related traits, especially under climate change conditions. In this research, grain quality, technological properties of flour, and some agronomic traits were studied in 33 wheat genotypes from six different subspecies of *Triticum turgidum* along with three cultivars of *Triticum aestivum* in the field, applying a well-watering (WW) and a water stress (WS) environment during two growing seasons. A high degree of variation was observed among genotypes for all evaluated traits, demonstrating that selection for these traits would be successful. Consequences of water stress were manifested as declined DM, TGW, GY, and LASRC; and significantly increased GPC, ZEL, GH, WAF, WSRC, SuSRC, and SCSRC compared to the well-watering condition. The reductions in the unextractable polymeric protein fraction and glutenin-to-gliadin ratio indicated a poorer grain yield quality, despite higher protein content. This study showed that the early-maturing genotypes had higher water absorption and pentosan, and therefore are more suitable for bread baking. In contrast, late-maturing genotypes are ideal for cookie and cracker production. Two subspecies of *T. turgidum* ssp. *durum* and *T. turgidum* ssp. *dicoccum* with high micronutrient densities and quality-related traits, and *T. turgidum* ssp. *oriental* due to having high values of grain protein content can be used to improve the quality of *T. aestivum* through cross-breeding programs. Based on the association of different traits with SRC values and other quality-related traits and PCA results, contrasting genotypes can be used to develop mapping populations for genome studies of grain quality and functional properties of flour in future studies.

## Introduction

Wheat is one of the most important staple-foods that supply about the 20% of the total calories and daily proteins worldwide [1]. This crop is also a noble source of vegetal protein, dietary fiber, calcium, zinc, fats, vitamins B-complex, carbohydrates, and energy than other cereals. Since the arable land area will not increase much beyond present levels, achieving high production through improved wheat yields is the demand of the 21st century to satisfy the increasing food demand of the growing population [2].

In arid and semiarid regions, global wheat production is considerably threatened by prolonged drought that is predicted to increase with progressive global climate change. The development of drought-tolerant wheat varieties is the ultimate mean of safeguarding the crop against the damaging effects of drought [3]. In this respect, determination of the genetic diversity existing within and between wheat populations remains the basis for elucidation of the genetic structure and improvement of different traits, including drought tolerance.

During domestication of wild emmer and later advancement of modern plant breeding, the genetic diversity of the cultivated wheat germplasm has been drastically narrowed, which could jeopardize future crop improvement. This genetic bottleneck severely eroded allelic variation in cultivated wheat and poses a potential threat of serious crop vulnerability to many adverse events, including global warming and climate changes [4]. The utilization of new diversity is essential to overcome the narrow genetic base in wheat. Local wheat cultivars or landraces with higher yield stability and better adaption to climate change represent an extensive reservoir of genetic variation for desirable traits such as tolerance to biotic and abiotic stresses [5].

The increasing attention to sustainable and organic agriculture and growing demand for high health-promoting foods has led to a renewed interest in ancient hulled wheat species as healthy grains among the different species of wheat. *Triticum dicoccum* is one of the ancient hulled wheat species globally and represents a valuable genetic resource to improve resistance to biotic and abiotic stresses in bread and durum wheat [6].

Grain quality of wheat is mainly the outcome of independent and interactive actions of several traits, including grain hardness, gluten protein quality, flour color, and starch quality. It is influenced by genetic and environmental conditions and biotic and abiotic stresses [7]. As a result, grain quality varies over a vast range from cultivar to cultivar, year to year, and across ecological environments. However, an insufficient supply of Zn and Fe is a widespread mineral malnutrition problem among the world’s population. Therefore, the development of micronutrient efficient genotypes requires identifying suitable germplasm with genetic potential and sufficient stability under different environments.

Evaluating flour quality is an essential task for breeders, millers, and bakers in selecting good-quality wheat cultivars with optimized performance [8]. There are many different test methods to evaluate various categories of wheat quality. Among them, Solvent Retention Capacity (SRC) is a unique diagnostic tool for predicting flour functionality, which examines the glutenin, gliadin, and pentosan (fibers) characteristics of the flour and the level of starch damage in the flour [9]. SRC is based on quantifying the enhanced swelling behavior of flour polymer networks in diagnostic solvents. It was developed to evaluate soft wheat flour functionality, but it has also been shown to assess the flour functionality of hard wheat products [8].

A unique feature of the present study is that it compares three genetic background sources, bread (*Triticum aestivum* L.), durum (*T. durum* L.), and emmer (*T. dicoccum* L.) wheat that can be exploited to accelerate the improvement of wheat for quality traits, especially under water stress condition. Our objectives in this study were to (*i*) determine the genetic variability and heritability of some quality-related traits in six different wheat subspecies under well-watering (WW) and water-stress (WS) conditions; and (ii) evaluate the association of phenological, agronomic, and quality-related traits to identify the most suitable combination of these traits through the indirect selection of promising genotypes for future breeding programs.

## Materials and methods

### Experimental site

This research was conducted during two years (2019 and 2020) at the research farm of the Isfahan University of Technology, located in Lavark, Najaf-Abad, Isfahan, Iran (32° 30′ N, 51° 20′ E, 1630m amsl) on a Typic Haplargid, silty clay loam soil, with pH 8.3. In this region, there is no rain during summer (from late May to mid-October); therefore, crops must be irrigated. Based on 40-year meteorological data, the region’s mean annual precipitation and temperature were 140 mm and 14.5 °C, respectively.

### Plant material and field evaluations

The genetic material used for this study consisted of thirty-six wheat genotypes, including thirty-three tetraploid genotypes that belonged to six different subspecies of *T. turgidum*, along with three bread wheat (*T. aestivum*) cultivars (Table 1). The experiment was arranged based on a randomized complete blocks design with four replications in two wheat cropping seasons.

**Table 1.** Information on genetic materials used in the study.

| Genotype | Public name or accession number | Ploidy level | Scientific name | Origin |
| --- | --- | --- | --- | --- |
| 1 | Shabrang | 4X | *T. turgidum* L. ssp. *durum* | Iran |
| 2 | Dena | 4X | *T. turgidum* L. ssp. *durum* | Iran |
| 3 | Aria | 4X | *T. turgidum* L. ssp. *durum* | Iran |
| 4 | Behrang | 4X | *T. turgidum* L. ssp. *durum* | Iran |
| 5 | Yavaros | 4X | *T. turgidum* L. ssp. *durum* | CIMMYT |
| 6 | Shwa | 4X | *T. turgidum* L. ssp. *durum* | ICARDA |
| 7 | Karkheh | 4X | *T. turgidum* L. ssp. *durum* | Turkey |
| 8 | Saji | 4X | *T. turgidum* L. ssp. *durum* | Iran |
| 9 | Khoygan | 4X | *T. turgidum* L. ssp. *dicoccum* | Iran |
| 10 | Ozonbelagh | 4X | *T. turgidum* L. ssp. *dicoccum* | Iran |
| 11 | Zarneh | 4X | *T. turgidum* L. ssp. *dicoccum* | Iran |
| 12 | Singerd | 4X | *T. turgidum* L. ssp. *dicoccum* | Iran |
| 13 | Berkman | 4X | *T. turgidum* L. ssp. *durum* | Turkey |
| 14 | 24 | 4X | *T. turgidum* L. ssp. *dicoccum* | Iran |
| 15 | 20 | 4X | *T. turgidum* L. ssp. *oreintal* | Iran |
| 16 | 6 | 4X | *T. turgidum* L. ssp. *oreintal* | Iran |
| 17 | 2 | 4X | *T. turgidum* L. ssp. *oreintal* | Iran |
| 18 | 19 | 4X | *T. turgidum* L. ssp. *turgidum* | Iran |
| 19 | 22 | 4X | *T. turgidum* L. ssp. *polonicum* | Iran |
| 20 | 13 | 4X | *T. turgidum* L. ssp. *polonicum* | Iran |
| 21 | 7 | 4X | *T. turgidum* L. ssp. *polonicum* | Iran |
| 22 | 3 | 4X | *T. turgidum* L. ssp. *polonicum* | Iran |
| 23 | 18 | 4X | *T. turgidum* L. ssp. *persicum* | Iran |
| 24 | 17 | 4X | *T. turgidum* L. ssp. *persicum* | Iran |
| 25 | 16 | 4X | *T. turgidum* L. ssp. *dicoccum* | Iran |
| 26 | 495 | 4X | *T. turgidum* L. ssp. *durum* | IPK |
| 27 | 493 | 4X | *T. turgidum* L. ssp. *durum* | IPK |
| 28 | Parsi | 6X | *T. aestivum* L. ssp. *aestivum* | Iran |
| 29 | 503 | 4X | *T. turgidum* L. ssp. *dicoccum* | IPK |
| 30 | 499 | 4X | *T. turgidum* L. ssp. *polonicum* | IPK |
| 31 | 496 | 4X | *T. turgidum* L. ssp. *persicum* | IPK |
| 32 | 502 | 4X | *T. turgidum* L. ssp. *turgidum* | IPK |
| 33 | 501/Z | 4X | *T. turgidum* L. ssp. *turgidum* | IPK |
| 34 | Golestan | 6X | *T. aestivum* L. ssp. *aestivum* | Iran |
| 35 | Tajan | 6X | *T. aestivum* L. ssp. *aestivum* | Iran |
| 36 | Zardak | 4X | *T. turgidum* L. ssp. *durum* | Iran |

In both years, the seeds of each genotype were sown in plots of two 100-cm long rows, 20-cm row spacing, and a within-row spacing of 2 cm, adjusting the seeding rate of 300 seeds m^−2^.

Plants were evaluated under WW and WS conditions. Two replications were allocated to each of these two moisture environments. Under the WW and WS conditions, water was supplied when 40 and 80% of plant available water (PAW) was exhausted from the root-zone, respectively. To determine the amount of irrigation water needed for supplying the soil moisture deficit to the field capacity and detect the irrigation times, soil samples were taken from different sites of each WW and WS environment every 2 days at depths of 0–20, 20–40, and 40–60 cm, using a hand auger and the gravimetric soil–water content was measured. At the irrigation time, the irrigation depth was determined as follows (Eq. 1):

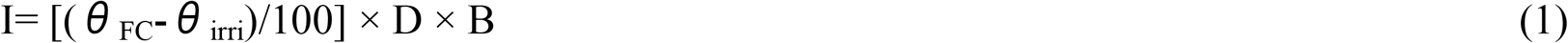

where I is the irrigation depth (cm), *θ*_FC_ is the soil gravimetric moisture percentage at field capacity, *θ*_irri_ is the soil gravimetric moisture percentage at irrigating time, D is the root-zone depth, and B is the soil bulk density at root-zone (1.4 g cm^−3^). At the second step, water volumes that should be applied in each moisture environment were calculated by multiplication of the irrigation depth and the total area of plots under each moisture environment. The depth of irrigation (I_g_) was calculated as follows (Eq. 2):

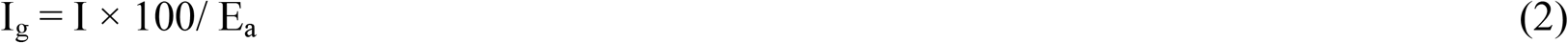

where I is the irrigation depth and E_a_ is the irrigation efficiency (%) assumed as 75% during the growing season. Water was delivered to the field using a drip irrigation system through a pumping station, polyethylene pipes, and drip tapes. The water volume applied under each moisture environment was measured using a volumetric counter.

Sixteen agro-morphological, quality, and SRC-related characteristics that were evaluated under the two levels of irrigation included: days to maturity (DM; days); grain yield (GY; g/m^2^); thousand grain weight (TGW; g); grain protein content (GPC; %); Zeleny index (ZEL; %); grain hardiness (GH; %); Zn (mg/g); Fe (mg/g); Na^+^ (mg/g); K^+^ (mg/g); moisture of flour (MOF; %); water absorption of flour (WAF; %); water SRC (WSRC; %); sucrose SRC (SuSRC; %); lactic acid SRC (LASRC; %); and sodium carbonate SRC (SCSRC; %). Five randomly selected samples of each genotype were used for trait measurement, and the mean value of five plants was calculated and used for analyses. Concentrations of Zn and Fe were determined by graphite furnace atomic absorption spectrometry (GFAAS) (PerkinElmer 800, PerkinElmer, Wellesley, MA). All grain concentrations are expressed on a dry weight basis. For determining the concentrations of Na^+^ and K^+^, grain samples (0.2 g) were ashed at 550 °C for 3 h. Inorganic ions were then extracted using 10 ml 2 N HCl, and the volume of each sample was standardized to 100 ml. Na^+^ and K^+^ concentrations of the solutions were determined by flame photometry (Jenway PFP7, UK). The amount of Na^+^ and K^+^ concentrations were calculated using a standard curve.

Solvent retention capacities (SRC) tests of the investigated genotypes were conducted according to the modified AACC International Approved Method 56-11.02 using 5% lactic acid, 50% sucrose, 5% sodium carbonate, and distilled water, with some modifications described by Duyvejonck et al. [10]. The amendment refers to the reduced mass of the sample. SRC is the weight of solvent held by flour after centrifugation; which is calculated as (eq. 3):

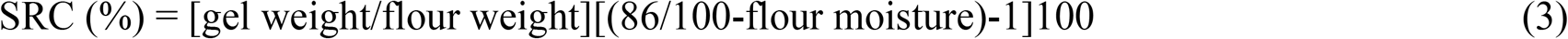

Each solvent diagnoses the functional contribution of specific flour components: lactic acid (LA) SRC is a measure for glutenin network forming capacity; sodium carbonate (SC) SRC is related to solvent-accessible amylopectin in damaged starch; sucrose (Su) SRC is a measure for the swelling of water-accessible pentosan (fiber); and finally, water SRC (WSRC) is related to overall water holding capacity by all network-forming components [8]. All solvents were obtained from Merck (KGaA 64271 Darmstadt, Germany). All SRC analyses were at least performed in duplicate, and the coefficient of variation of the SRC values was less than 5.0%.

### Statistical analyses

Before analysis of variance (ANOVA), the Kolmogorov–Smirnov test was conducted to examine the normality distribution of data. The Bartlett test was used to test the homogeneity of residual variance. Subsequently, to examine the differences between the genotypes, years, moisture environments, and all possible interactions, and also to estimate the variance components, combined analysis of variance was conducted using Proc MIXED of SAS release 9.4 (SAS Institute, Cary, NC, USA). The genotype effect was considered fixed, and year was considered random effect. Where the *F*-value was significant, mean comparisons were carried out using the least significant difference (LSD) test at *p* ˂ .05 [11]. Phenotypic correlation coefficients between traits were calculated using proc CORR of SAS to determine the association between evaluated characteristics. Broad-sense heritability (h^2^_b_) was estimated on a phenotypic mean basis averaged over replications, years and environments according to the following formula (Eq. 4):

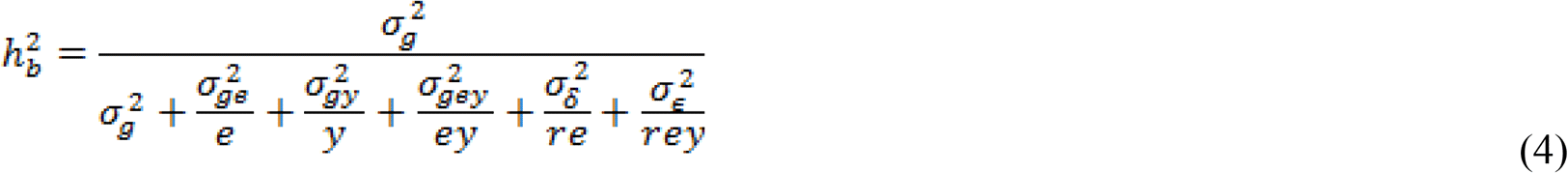

where h^2^_b_ is the broad-sense heritability, 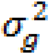 is the genotype, 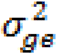 is the genotype × environment, 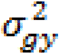 is the genotype × year, 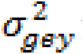 is the genotype × environment × year variance; 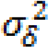 and 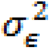 are the error variance and the residual variance, respectively; while g, e, y, and r represent the number of genotypes, environments, years, and replications, respectively.

The level of genetic variation was estimated with the calculation of phenotypic coefficient of variation (PCV) and genotypic coefficient of variation (GCV) as follows (Eqs. 5 and 6):

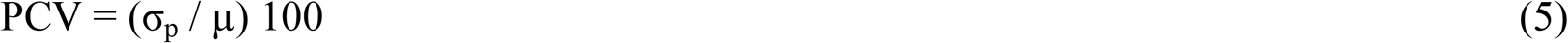

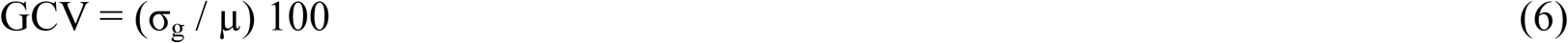

where σ_p_ is the standard deviation of the phenotypic variance, σ_g_ is the standard deviation of the genotypic variance, and μ is the phenotypic mean [12]. Data were also subjected to ANOVA separately for WW and WS across years. Variance components were estimated for individual moisture environments (WW and WS environments) using proc MIXED of SAS. Broad-sense heritability on a phenotypic mean basis averaged over replications and years was estimated as (Eq. 7):

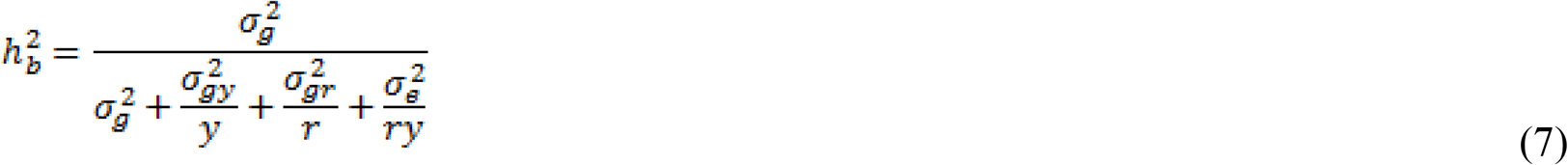

where h^2^_b_ is the broad-sense heritability, 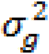 is the genotype, 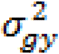 is the genotype × year, 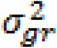 is the genotype × replication variance; and 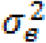 is the error variance; while g, y, and r represent the number of genotypes, years, and replications, respectively. Principal component analysis (PCA) was performed based on a correlation matrix by using Statgraphics software version 17.2 (Statgraphics Technologies, Inc.).

## Results

Combined analysis of variance indicated that there were significant differences (P ˂ 0.05) between the WW and WS environments for all traits except for ZEL. The effect of genotype was significant (P ˂ 0.01) for all of the evaluated traits; indicating considerable genotypic variation among the selected genotypes with a broad range for each attribute (Tables 2 and S1). Differentiation of genotype effect into subspecies showed non-significant genetic diversity for most studied traits and subspecies (Tables 2 and 3). Genotype × environment (GE) interaction was significant for DM, GY, Na^+^, and SCSRC; which show the different response of genotypes to environmental variations in terms of these traits. Contrastingly, genotypes showed similar responses across the two conditions for the remaining traits (Tables 2 and S1).

**Table 2.** Combined analysis of variance for measured traits in 36 wheat genotypes evaluated under two moisture environments (well-watering and water stress) during two years (2019 and 2020).

| Traits | df | Mean squares |  |  |  |  |  |  |  |
| --- | --- | --- | --- | --- | --- | --- | --- | --- | --- |
|  |  | DM | TGW | GY | PGP | ZEL | GH | Zn | Fe |
| Environment (E) | 1 | 3997.67** | 1059.26** | 1222423.78** | 86.61** | 36.46<sup>n.s</sup> | 380.42 * | 0.101 * | 0.204 * |
| Year (Y) | 1 | 16095.17** | 840.67* | 162.16<sup>n.s</sup> | 63.39** | 0.93<sup>n.s</sup> | 4057.50<sup>n.s</sup> | 0.023<sup>n.s</sup> | 0.006<sup>n.s</sup> |
| Y × E | 1 | 25.09<sup>n.s</sup> | 48.55<sup>n.s</sup> | 8224.14<sup>n.s</sup> | 0.10<sup>n.s</sup> | 15.21<sup>n.s</sup> | 509.34 ** | 0.279 ** | 0.103<sup>n.s</sup> |
| Replication (Y × E) | 4 | 26.79** | 99.22** | 23697.61<sup>n.s</sup> | 0.44<sup>n.s</sup> | 1.74<sup>n.s</sup> | 20.91<sup>n.s</sup> | 0.004<sup>n.s</sup> | 0.043 ** |
| Genotype (G) | 35 | 266.72** | 305.34** | 53699.34** | 11.15** | 14.84 ** | 243.76 ** | 0.027 ** | 0.031 ** |
| *T. durum* | 11 | 54.00* | 172.59* | 93705.33* | 3.76* | 20.89 * | 290.31 ** | 0.010 ** | 0.012<sup>n.s</sup> |
| *T. dicoccum* | 6 | 12.89<sup>n.s</sup> | 45.55<sup>n.s</sup> | 44075.60* | 3.96<sup>n.s</sup> | 3.89<sup>n.s</sup> | 12.20 * | 0.022 ** | 0.017 ** |
| *T. oriental* | 2 | 9.54<sup>n.s</sup> | 0.89<sup>n.s</sup> | 1464.76<sup>n.s</sup> | 0.02<sup>n.s</sup> | 0.27<sup>n.s</sup> | 0.54<sup>n.s</sup> | 0.001<sup>n.s</sup> | 0.002<sup>n.s</sup> |
| *T. turgidum* | 2 | 1.17<sup>n.s</sup> | 455.63<sup>n.s</sup> | 5343.86<sup>n.s</sup> | 1.58<sup>n.s</sup> | 1.74<sup>n.s</sup> | 356.79 ** | 0.016 * | 0.006<sup>n.s</sup> |
| *T. polonicum* | 4 | 69.03<sup>n.s</sup> | 473.55** | 148925.20* | 2.04<sup>n.s</sup> | 3.56<sup>n.s</sup> | 143.04<sup>n.s</sup> | 0.032 * | 0.026<sup>n.s</sup> |
| *T. persicum* | 2 | 69.13<sup>n.s</sup> | 45.14<sup>n.s</sup> | 235046.62* | 6.09<sup>n.s</sup> | 25.11 ** | 0.67<sup>n.s</sup> | 0.002<sup>n.s</sup> | 0.029 * |
| *T. aestivum* | 2 | 28.04<sup>n.s</sup> |  | 30765.41<sup>n.s</sup> | 2.93<sup>n.s</sup> | 11.83<sup>n.s</sup> | 242.79 ** | 0.009<sup>n.s</sup> | 0.004<sup>n.s</sup> |
|  |  |  | 120.60<sup>n.s</sup> |  |  |  |  |  |  |
| G × E | 35 | 27.23** | 14.80<sup>n.s</sup> | 28413.17** | 0.72<sup>n.s</sup> | 8.94<sup>n.s</sup> | 20.43<sup>n.s</sup> | 0.007<sup>n.s</sup> | 0.009<sup>n.s</sup> |
| Y × G | 35 | 30.94** | 61.11** | 12845.85<sup>n.s</sup> | 0.91<sup>n.s</sup> | 10.36<sup>n.s</sup> | 37.28<sup>n.s</sup> | 0.009<sup>n.s</sup> | 0.012<sup>n.s</sup> |
| Y × G × E | 35 | 5.40<sup>n.s</sup> | 21.91<sup>n.s</sup> | 13405.08<sup>n.s</sup> | 0.71** | 10.57 ** | 40.93 ** | 0.006<sup>n.s</sup> | 0.009<sup>n.s</sup> |
| Error | 140 | 7.68 | 17.90 | 12462.11 | 0.18 | 5.90 | 17.86 | 0.004 | 0.008 |

Table 2- Continued
| Traits | df | Mean squares |  |  |  |  |  |  |  |
| --- | --- | --- | --- | --- | --- | --- | --- | --- | --- |
|  |  | Na<sup>+</sup> | K<sup>+</sup> | MOF | WAF | WSRC | SuSRC | LASRC | SCSRC |
| Environment (E) | 1 | 1.68 * | 41.69 * | 18.50 * | 41.25 * | 3370.24 ** | 3585.08 * | 2427.58 * | 2866.85 ** |
| Year (Y) | 1 | 5.93 * | 3.15 n.s | 839.82 ** | 227.56 n.s | 5339.60 n.s | 9114.46 ** | 22234.18 ** | 27.58 n.s |
| Y × E | 1 | 0.03 n.s | 1.48 n.s | 0.12 n.s | 7.41 * | 840.56 n.s | 2.83 n.s | 282.67 n.s | 20.91 n.s |
| Replication (Y × E) | 4 | 0.81 ** | 2.84 ** | 0.42 ** | 0.17 n.s | 70.19 ** | 51.76 n.s | 164.34 ** | 68.30 n.s |
| Genotype (G) | 35 | 3.96 ** | 2.12 ** | 0.50 ** | 25.21 ** | 959.85 ** | 1088.57 ** | 2216.39 ** | 996.34 ** |
| *T. durum* | 11 | 5.85 ** | 2.78 ** | 0.22 * | 22.08 ** | 719.97 n.s | 877.30 n.s | 2133.73 * | 1080.57 * |
| *T. dicoccum* | 6 | 1.03 ** | 1.54 * | 0.15 n.s | 1.42 n.s | 103.57 n.s | 262.44 n.s | 271.36 n.s | 581.74 n.s |
| *T. oriental* | 2 | 0.01 n.s | 0.28 n.s | 0.03 n.s | 0.14 n.s | 14.70 n.s | 18.74 n.s | 104.56 n.s | 534.84 n.s |
| *T. turgidum* | 2 | 0.68 * | 0.50 n.s | 0.33 ** | 19.89 ** | 5.07 n.s | 38.12 n.s | 525.94 ** | 84.04 n.s |
| *T. polonicum* | 4 | 1.01 ** | 2.63 n.s | 0.14 * | 1.56 * | 286.76 n.s | 475.28 n.s | 637.47 n.s | 498.23 n.s |
| *T. persicum* | 2 | 5.07 ** | 2.49 * | 0.30 n.s | 0.59 n.s | 2281.19 n.s | 1773.38 n.s | 4079.74 n.s | 3068.47 n.s |
| *T. aestivum* | 2 | 0.01 n.s | 0.87 n.s | 0.06 n.s | 21.25 ** | 710.74 ** | 1086.17 ** | 1547.23 ** | 706.78 ** |
| G × E | 35 | 0.71 * | 0.93 n.s | 0.15 n.s | 1.85 n.s | 260.84 n.s | 219.36 n.s | 232.96 n.s | 247.46 * |
| Y × G | 35 | 1.59 ** | 1.36 n.s | 0.42 * | 1.66 n.s | 271.90 n.s | 334.54 * | 529.45 ** | 156.04 n.s |
| Y × G × E | 35 | 0.41 ** | 1.16 ** | 0.24 ** | 1.40 ** | 175.15 ** | 158.01 ** | 176.38 ** | 123.51 ** |
| Error | 140 | 0.14 | 0.50 | 0.07 | 0.50 | 14.42 | 26.24 | 29.79 | 29.21 |

\* and \*\* show significance at the 0.05 and 0.01 probability levels, respectively.
n.s: not significant
DM, days to maturity; Fe, Iron content; GH, grain hardness; GPC, grain protein content; GY, grain yield; K<sup>+</sup>, potassium content; LASRC, lactic acid SRC; MOF, moisture of flour; Na<sup>+</sup>, sodium content; SCSRC, sodium carbonate SRC; SuSRC, sucrose SRC; TGW, thousand grain weight; WAF, water absorption of flour; WSRC, water SRC; ZEL, Zeleny index; Zn, zinc content.

**Table 3.**
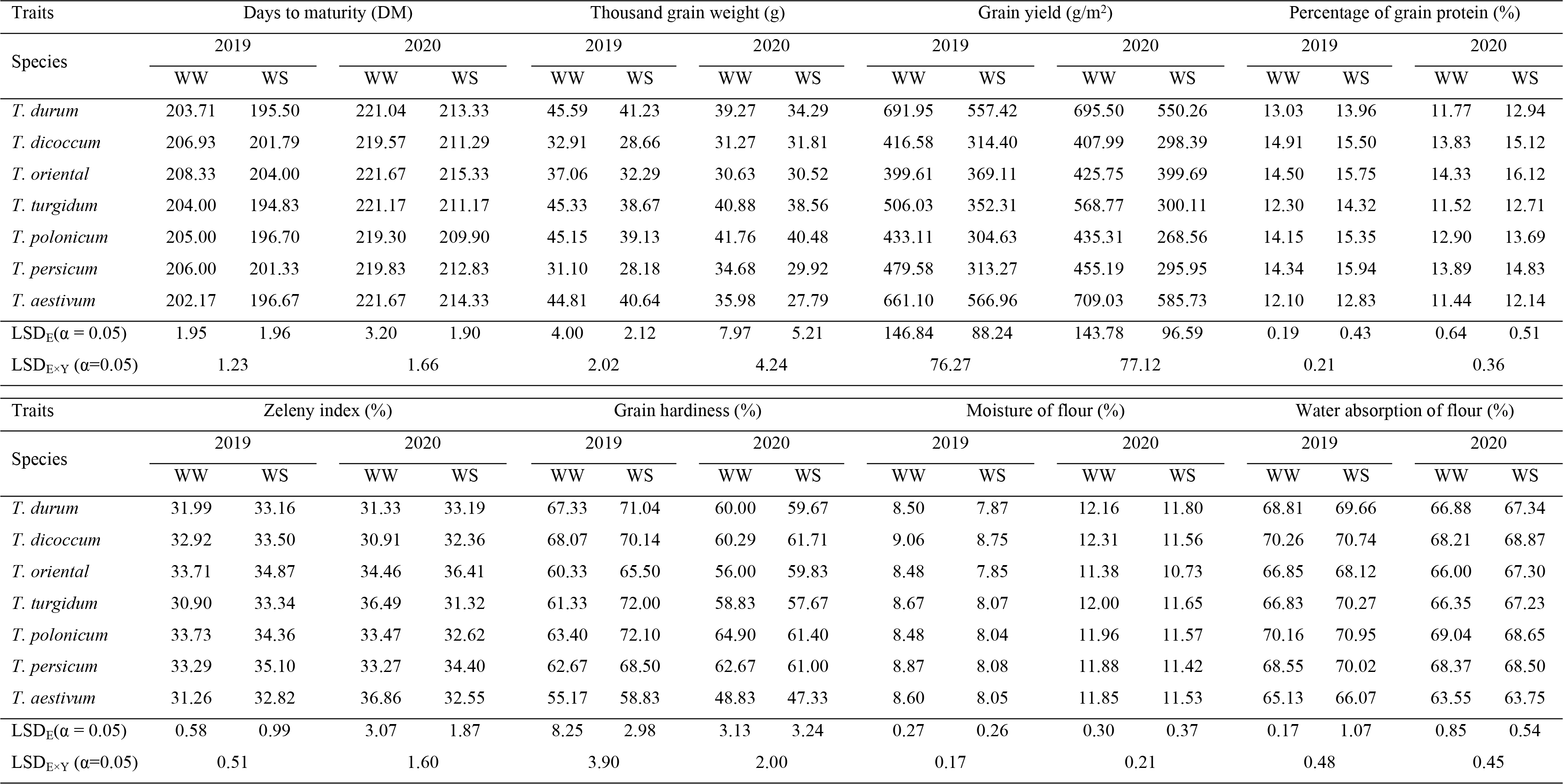
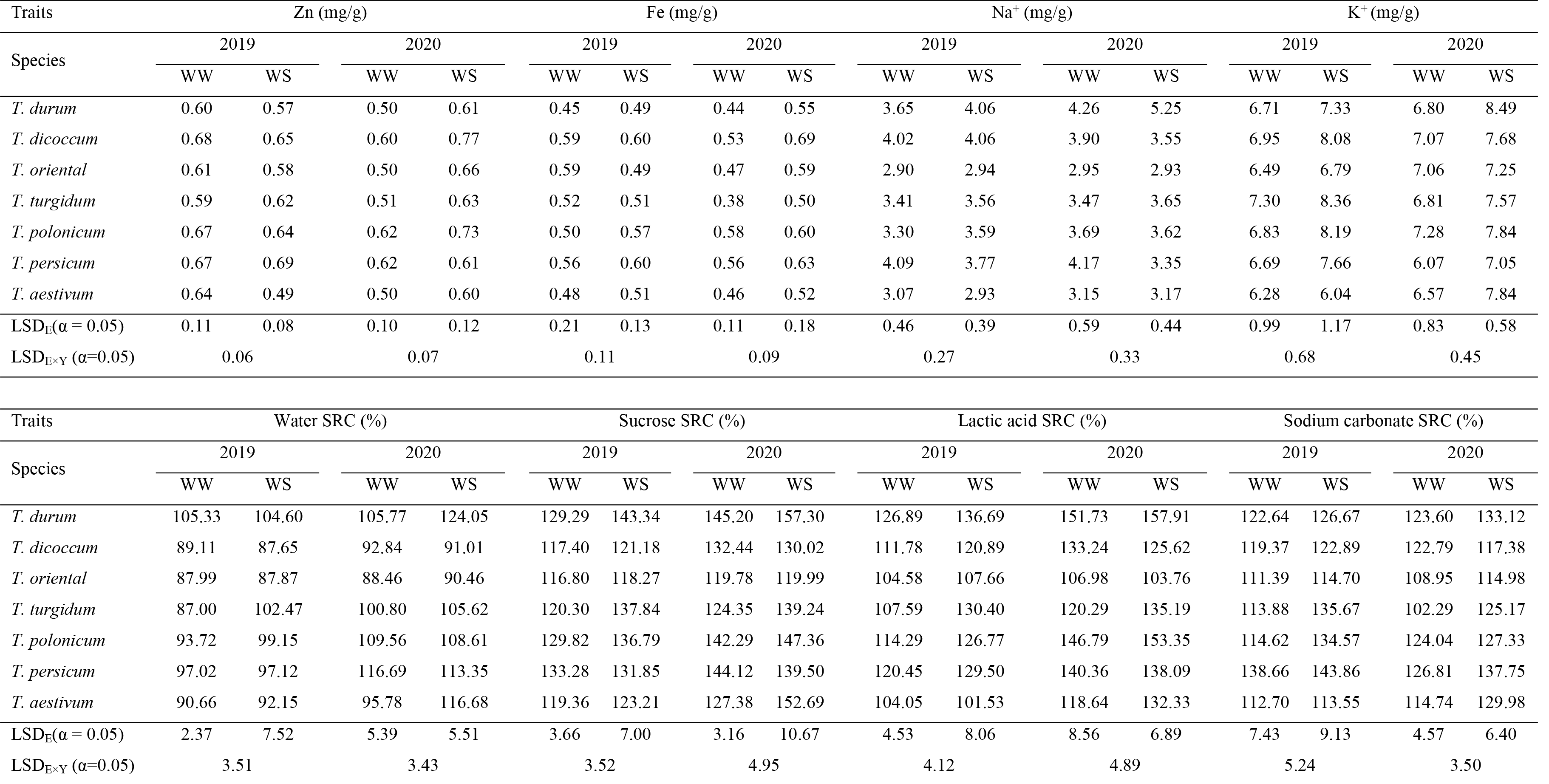
Mean comparisons of different traits evaluated on six tetraploid sub-species of wheat and Triticum aestivum during two years (2019 and 2020) under well-watering (WW) and water stress (WS) conditions.

| Traits | Days to maturity (DM) |  |  |  | Thousand grain weight (g) |  |  |  | Grain yield (g/m<sup>2</sup>) |  |  |  | Percentage of grain protein (%) |  |  |  |
| --- | --- | --- | --- | --- | --- | --- | --- | --- | --- | --- | --- | --- | --- | --- | --- | --- |
| Species | 2019 |  | 2020 |  | 2019 |  | 2020 |  | 2019 |  | 2020 |  | 2019 |  | 2020 |  |
|  | WW | WS | WW | WS | WW | WS | WW | WS | WW | WS | WW | WS | WW | WS | WW | WS |
| *T. durum* | 203.71 | 195.50 | 221.04 | 213.33 | 45.59 | 41.23 | 39.27 | 34.29 | 691.95 | 557.42 | 695.50 | 550.26 | 13.03 | 13.96 | 11.77 | 12.94 |
| *T. dicoccum* | 206.93 | 201.79 | 219.57 | 211.29 | 32.91 | 28.66 | 31.27 | 31.81 | 416.58 | 314.40 | 407.99 | 298.39 | 14.91 | 15.50 | 13.83 | 15.12 |
| *T. oriental* | 208.33 | 204.00 | 221.67 | 215.33 | 37.06 | 32.29 | 30.63 | 30.52 | 399.61 | 369.11 | 425.75 | 399.69 | 14.50 | 15.75 | 14.33 | 16.12 |
| *T. turgidum* | 204.00 | 194.83 | 221.17 | 211.17 | 45.33 | 38.67 | 40.88 | 38.56 | 506.03 | 352.31 | 568.77 | 300.11 | 12.30 | 14.32 | 11.52 | 12.71 |
| *T. polonicum* | 205.00 | 196.70 | 219.30 | 209.90 | 45.15 | 39.13 | 41.76 | 40.48 | 433.11 | 304.63 | 435.31 | 268.56 | 14.15 | 15.35 | 12.90 | 13.69 |
| *T. persicum* | 206.00 | 201.33 | 219.83 | 212.83 | 31.10 | 28.18 | 34.68 | 29.92 | 479.58 | 313.27 | 455.19 | 295.95 | 14.34 | 15.94 | 13.89 | 14.83 |
| *T. aestivum* | 202.17 | 196.67 | 221.67 | 214.33 | 44.81 | 40.64 | 35.98 | 27.79 | 661.10 | 566.96 | 709.03 | 585.73 | 12.10 | 12.83 | 11.44 | 12.14 |
| LSD<sub>E</sub>($\alpha = 0.05$) | 1.95 | 1.96 | 3.20 | 1.90 | 4.00 | 2.12 | 7.97 | 5.21 | 146.84 | 88.24 | 143.78 | 96.59 | 0.19 | 0.43 | 0.64 | 0.51 |
| LSD<sub>E×Y</sub> ($\alpha=0.05$) | 1.23 | | 1.66 | | 2.02 | | 4.24 | | 76.27 | | 77.12 | | 0.21 | | 0.36 | |

| Traits | Zeleny index (%) |  |  |  | Grain hardness (%) |  |  |  | Moisture of flour (%) |  |  |  | Water absorption of flour (%) |  |  |  |
| --- | --- | --- | --- | --- | --- | --- | --- | --- | --- | --- | --- | --- | --- | --- | --- | --- |
| Species | 2019 |  | 2020 |  | 2019 |  | 2020 |  | 2019 |  | 2020 |  | 2019 |  | 2020 |  |
|  | WW | WS | WW | WS | WW | WS | WW | WS | WW | WS | WW | WS | WW | WS | WW | WS |
| *T. durum* | 31.99 | 33.16 | 31.33 | 33.19 | 67.33 | 71.04 | 60.00 | 59.67 | 8.50 | 7.87 | 12.16 | 11.80 | 68.81 | 69.66 | 66.88 | 67.34 |
| *T. dicoccum* | 32.92 | 33.50 | 30.91 | 32.36 | 68.07 | 70.14 | 60.29 | 61.71 | 9.06 | 8.75 | 12.31 | 11.56 | 70.26 | 70.74 | 68.21 | 68.87 |
| *T. oriental* | 33.71 | 34.87 | 34.46 | 36.41 | 60.33 | 65.50 | 56.00 | 59.83 | 8.48 | 7.85 | 11.38 | 10.73 | 66.85 | 68.12 | 66.00 | 67.30 |
| *T. turgidum* | 30.90 | 33.34 | 36.49 | 31.32 | 61.33 | 72.00 | 58.83 | 57.67 | 8.67 | 8.07 | 12.00 | 11.65 | 66.83 | 70.27 | 66.35 | 67.23 |
| *T. polonicum* | 33.73 | 34.36 | 33.47 | 32.62 | 63.40 | 72.10 | 64.90 | 61.40 | 8.48 | 8.04 | 11.96 | 11.57 | 70.16 | 70.95 | 69.04 | 68.65 |
| *T. persicum* | 33.29 | 35.10 | 33.27 | 34.40 | 62.67 | 68.50 | 62.67 | 61.00 | 8.87 | 8.08 | 11.88 | 11.42 | 68.55 | 70.02 | 68.37 | 68.50 |
| *T. aestivum* | 31.26 | 32.82 | 36.86 | 32.55 | 55.17 | 58.83 | 48.83 | 47.33 | 8.60 | 8.05 | 11.85 | 11.53 | 65.13 | 66.07 | 63.55 | 63.75 |
| LSD<sub>E</sub>($\alpha = 0.05$) | 0.58 | 0.99 | 3.07 | 1.87 | 8.25 | 2.98 | 3.13 | 3.24 | 0.27 | 0.26 | 0.30 | 0.37 | 0.17 | 1.07 | 0.85 | 0.54 |
| LSD<sub>E×Y</sub> ($\alpha=0.05$) | 0.51 | | 1.60 | | 3.90 | | 2.00 | | 0.17 | | 0.21 | | 0.48 | | 0.45 | |

**Table 3-** Continued
| Traits | Zn (mg/g) |  |  |  | Fe (mg/g) |  |  |  | Na<sup>+</sup> (mg/g) |  |  |  | K<sup>+</sup> (mg/g) |  |  |  |
| --- | --- | --- | --- | --- | --- | --- | --- | --- | --- | --- | --- | --- | --- | --- | --- | --- |
| Species | 2019 |  | 2020 |  | 2019 |  | 2020 |  | 2019 |  | 2020 |  | 2019 |  | 2020 |  |
|  | WW | WS | WW | WS | WW | WS | WW | WS | WW | WS | WW | WS | WW | WS | WW | WS |
| *T. durum* | 0.60 | 0.57 | 0.50 | 0.61 | 0.45 | 0.49 | 0.44 | 0.55 | 3.65 | 4.06 | 4.26 | 5.25 | 6.71 | 7.33 | 6.80 | 8.49 |
| *T. dicoccum* | 0.68 | 0.65 | 0.60 | 0.77 | 0.59 | 0.60 | 0.53 | 0.69 | 4.02 | 4.06 | 3.90 | 3.55 | 6.95 | 8.08 | 7.07 | 7.68 |
| *T. oriental* | 0.61 | 0.58 | 0.50 | 0.66 | 0.59 | 0.49 | 0.47 | 0.59 | 2.90 | 2.94 | 2.95 | 2.93 | 6.49 | 6.79 | 7.06 | 7.25 |
| *T. turgidum* | 0.59 | 0.62 | 0.51 | 0.63 | 0.52 | 0.51 | 0.38 | 0.50 | 3.41 | 3.56 | 3.47 | 3.65 | 7.30 | 8.36 | 6.81 | 7.57 |
| *T. polonicum* | 0.67 | 0.64 | 0.62 | 0.73 | 0.50 | 0.57 | 0.58 | 0.60 | 3.30 | 3.59 | 3.69 | 3.62 | 6.83 | 8.19 | 7.28 | 7.84 |
| *T. persicum* | 0.67 | 0.69 | 0.62 | 0.61 | 0.56 | 0.60 | 0.56 | 0.63 | 4.09 | 3.77 | 4.17 | 3.35 | 6.69 | 7.66 | 6.07 | 7.05 |
| *T. aestivum* | 0.64 | 0.49 | 0.50 | 0.60 | 0.48 | 0.51 | 0.46 | 0.52 | 3.07 | 2.93 | 3.15 | 3.17 | 6.28 | 6.04 | 6.57 | 7.84 |
| LSD<sub>E</sub>($\alpha = 0.05$) | 0.11 | 0.08 | 0.10 | 0.12 | 0.21 | 0.13 | 0.11 | 0.18 | 0.46 | 0.39 | 0.59 | 0.44 | 0.99 | 1.17 | 0.83 | 0.58 |
| LSD<sub>E×Y</sub> ($\alpha=0.05$) | 0.06 | | 0.07 | | 0.11 | | 0.09 | | 0.27 | | 0.33 | | 0.68 | | 0.45 | |

| Traits | Water SRC (%) |  |  |  | Sucrose SRC (%) |  |  |  | Lactic acid SRC (%) |  |  |  | Sodium carbonate SRC (%) |  |  |  |
| --- | --- | --- | --- | --- | --- | --- | --- | --- | --- | --- | --- | --- | --- | --- | --- | --- |
| Species | 2019 |  | 2020 |  | 2019 |  | 2020 |  | 2019 |  | 2020 |  | 2019 |  | 2020 |  |
|  | WW | WS | WW | WS | WW | WS | WW | WS | WW | WS | WW | WS | WW | WS | WW | WS |
| *T. durum* | 105.33 | 104.60 | 105.77 | 124.05 | 129.29 | 143.34 | 145.20 | 157.30 | 126.89 | 136.69 | 151.73 | 157.91 | 122.64 | 126.67 | 123.60 | 133.12 |
| *T. dicoccum* | 89.11 | 87.65 | 92.84 | 91.01 | 117.40 | 121.18 | 132.44 | 130.02 | 111.78 | 120.89 | 133.24 | 125.62 | 119.37 | 122.89 | 122.79 | 117.38 |
| *T. oriental* | 87.99 | 87.87 | 88.46 | 90.46 | 116.80 | 118.27 | 119.78 | 119.99 | 104.58 | 107.66 | 106.98 | 103.76 | 111.39 | 114.70 | 108.95 | 114.98 |
| *T. turgidum* | 87.00 | 102.47 | 100.80 | 105.62 | 120.30 | 137.84 | 124.35 | 139.24 | 107.59 | 130.40 | 120.29 | 135.19 | 113.88 | 135.67 | 102.29 | 125.17 |
| *T. polonicum* | 93.72 | 99.15 | 109.56 | 108.61 | 129.82 | 136.79 | 142.29 | 147.36 | 114.29 | 126.77 | 146.79 | 153.35 | 114.62 | 134.57 | 124.04 | 127.33 |
| *T. persicum* | 97.02 | 97.12 | 116.69 | 113.35 | 133.28 | 131.85 | 144.12 | 139.50 | 120.45 | 129.50 | 140.36 | 138.09 | 138.66 | 143.86 | 126.81 | 137.75 |
| *T. aestivum* | 90.66 | 92.15 | 95.78 | 116.68 | 119.36 | 123.21 | 127.38 | 152.69 | 104.05 | 101.53 | 118.64 | 132.33 | 112.70 | 113.55 | 114.74 | 129.98 |
| LSD<sub>E</sub>($\alpha = 0.05$) | 2.37 | 7.52 | 5.39 | 5.51 | 3.66 | 7.00 | 3.16 | 10.67 | 4.53 | 8.06 | 8.56 | 6.89 | 7.43 | 9.13 | 4.57 | 6.40 |
| LSD<sub>E×Y</sub> ($\alpha=0.05$) | 3.51 | | 3.43 | | 3.52 | | 4.95 | | 4.12 | | 4.89 | | 5.24 | | 3.50 | |

Results of this study showed that WS significantly decreased DM, GY, and LASRC; while significantly increased GPC, WAF, WSRC, SuSRC, and SCSRC in both years of the study. Moreover, TGW significantly decreased and ZEL increased due to WS in 2019, but they were not influenced during 2020. During 2020, Zn significantly increased, and MOF decreased under WS condition; while they were not affected in 2019 (Table 4). The magnitude of mean performance of the remaining traits was not affected under WS condition. As expected, WS reduced grain yield on average by 22 and 26% in 2019 and 2020, respectively. However, GPC increased approximately by 15 and 9% in the two consecutive years under WS condition (Table 4).

**Table 4.** The effect of water stress, phenotypic coefficient of variation (PCV), and genotypic coefficient of variation (GCV) for different traits recorded under well-watering (WW) and water stress (WS) conditions in wheat during two years (2019 and 2020).

| Traits | 2019 |  |  | 2020 |  |  | PCV |  | GCV |  |
| --- | --- | --- | --- | --- | --- | --- | --- | --- | --- | --- |
|  | WW | WS | Change (%) | WW | WS | Change (%) | WW | WS | WW | WS |
| DM (day) | 204.99 | 198.13 | -3.35 ** | 220.53 | 212.49 | -3.65 * | 2.04 | 1.87 | 1.84 | 1.47 |
| TGW (g) | 41.06 | 36.40 | -11.35 * | 36.82 | 33.81 | -8.17 n.s | 17.59 | 16.39 | 15.76 | 13.24 |
| GY (g/m<sup>2</sup>) | 542.33 | 422.72 | -22.05 * | 551.52 | 410.53 | -25.56 * | 9.00 | 19.51 | 6.99 | 14.21 |
| GPC (%) | 13.64 | 14.70 | 7.77 * | 12.67 | 13.80 | 8.92 ** | 9.13 | 8.66 | 8.78 | 7.77 |
| ZEL (%) | 32.51 | 33.68 | 3.60 * | 32.86 | 33.11 | 0.76 n.s | 8.27 | 5.50 | 6.00 | 3.80 |
| GH (%) | 64.44 | 69.40 | 7.70 n.s | 59.60 | 59.24 | -0.60 n.s | 9.56 | 8.64 | 7.55 | 7.71 |
| Zn (mg/g) | 0.63 | 0.60 | -4.76 n.s | 0.55 | 0.66 | 20.00 ** | 10.89 | 11.30 | 9.64 | 7.56 |
| Fe (mg/g) | 0.51 | 0.53 | 3.92 n.s | 0.50 | 0.58 | 16.00 n.s | 14.72 | 12.69 | 9.62 | 9.65 |
| Na<sup>+</sup> (mg/g) | 3.59 | 3.75 | 4.46 n.s | 3.89 | 4.04 | 3.86 n.s | 17.09 | 22.87 | 12.91 | 17.05 |
| K<sup>+</sup> (mg/g) | 6.76 | 7.54 | 11.54 n.s | 6.91 | 7.89 | 14.18 n.s | 9.78 | 11.66 | 7.41 | 7.93 |
| MOF (%) | 8.66 | 8.11 | -6.35 n.s | 12.03 | 11.57 | -3.82 * | 2.53 | 4.18 | 1.49 | 2.26 |
| WAF (%) | 68.63 | 69.70 | 1.56 ** | 67.17 | 67.60 | 0.64 * | 2.77 | 2.29 | 2.62 | 2.10 |
| WSRC (%) | 95.68 | 107.87 | 12.74 ** | 102.01 | 119.64 | 17.28 ** | 11.06 | 13.06 | 8.87 | 10.26 |
| SuSRC (%) | 124.76 | 143.72 | 15.20 ** | 138.07 | 154.13 | 11.63 ** | 8.30 | 10.45 | 6.65 | 8.12 |
| LASRC (%) | 136.30 | 116.45 | -14.56 ** | 149.61 | 130.81 | -12.57 ** | 12.72 | 14.09 | 10.46 | 11.92 |
| SCSRC (%) | 119.73 | 137.62 | 14.94 ** | 121.31 | 137.20 | 13.10 ** | 9.02 | 11.00 | 7.97 | 9.69 |

DM, days to maturity; Fe, Iron content; GH, grain hardness; GPC, grain protein content; GY, grain yield; K<sup>+</sup>, potassium content; LASRC, lactic acid SRC; MOF, moisture of flour; Na<sup>+</sup>, sodium content; SCSRC, sodium carbonate SRC; SuSRC, sucrose SRC; TGW, thousand grain weight; WAF, water absorption of flour; WSRC, water SRC; ZEL, Zeleny index; Zn, zinc content.

Results also revealed that, among the studied subspecies *T. polonicum* was the earliest maturing subspecies and had the highest TGW, GH, Zn, K^+^, and WAF; and along with the subspecies of *T. dicoccum* showed the lowest value of GY. The subspecies of *T. oriental* was the latest one and showed the highest values of GPC and ZEL index, and had the lowest values in terms of Na^+^, MOF, WSRC, SuSRC, LASRC, and SCSRC. Moreover, *T. durum* showed the highest GY, Na^+^, WSRC, SuSRC, and LASRC, and the lowest ZEL, Zn and Fe. The highest values of GH, Zn, Fe, MOF, and WAF, and the lowest values of GY and ZEL were allocated to *T. dicoccum*. *Triticum aestivum* showed the highest grain yield and the lowest values of GPC, GH, Zn, Fe, K^+^, and WAF. *T. persicum* had the highest SCSRC and the lowest value of TGW; while the other traits showed intermediate values in this subspecies. *T. turgidum* had the highest value of K^+^ and the lowest value of Fe; while other traits showed intermediate values in it (Table 5).

**Table 5.**
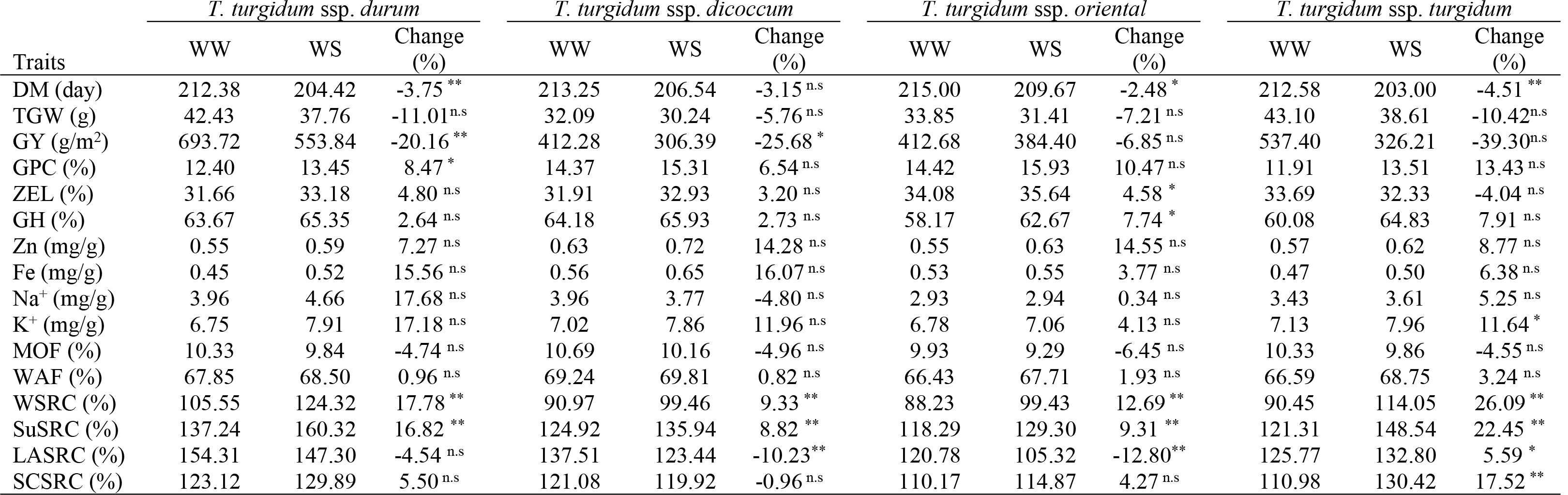
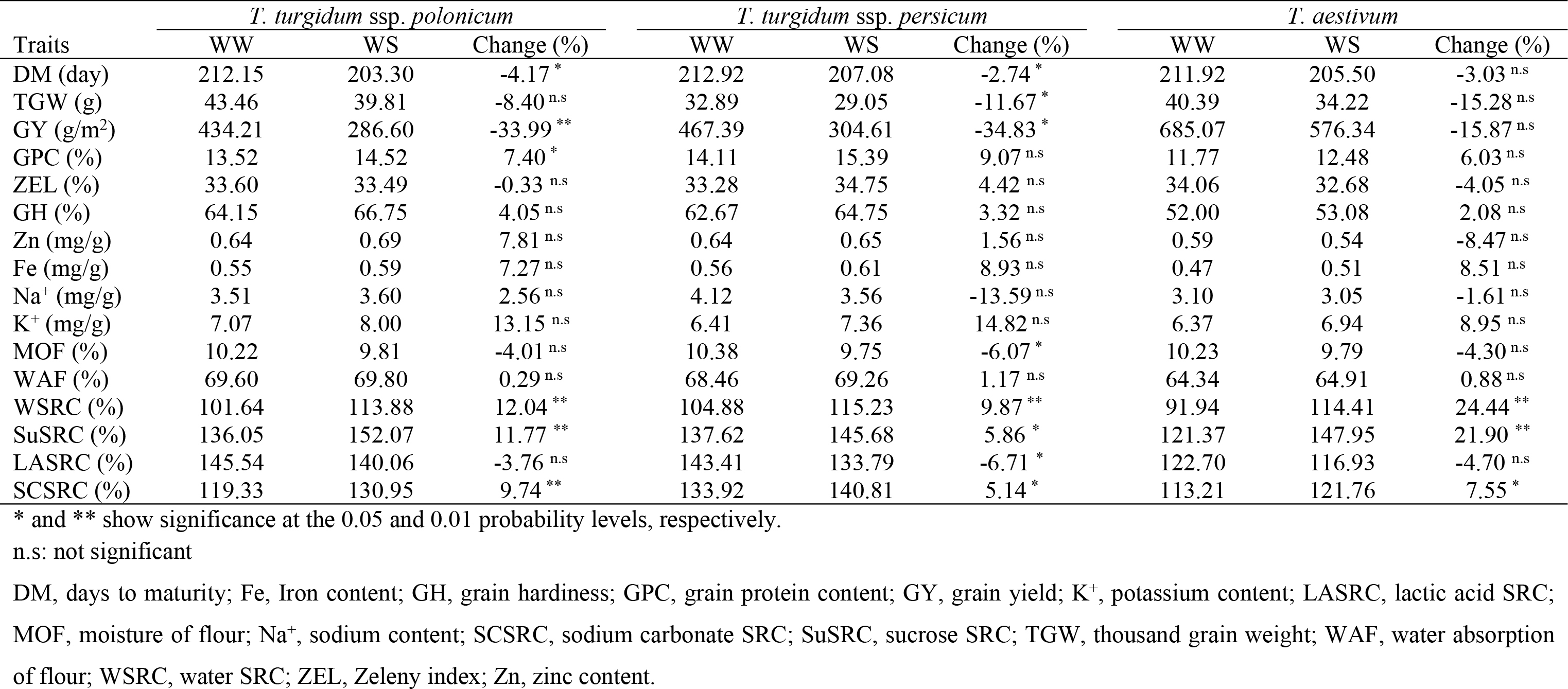
The effect of water stress on different traits evaluated under well-watering (WW) and water stress (WS) conditions in seven subspecies of wheat during two years (2019 and 2020).

| Traits | *T. turgidum* ssp. *durum* |  |  | *T. turgidum* ssp. *dicoccum* |  |  | *T. turgidum* ssp. *oriental* |  |  | *T. turgidum* ssp. *turgidum* |  |  |
| --- | --- | --- | --- | --- | --- | --- | --- | --- | --- | --- | --- | --- |
|  | WW | WS | Change (%) | WW | WS | Change (%) | WW | WS | Change (%) | WW | WS | Change (%) |
| DM (day) | 212.38 | 204.42 | -3.75 ** | 213.25 | 206.54 | -3.15 n.s | 215.00 | 209.67 | -2.48 * | 212.58 | 203.00 | -4.51 ** |
| TGW (g) | 42.43 | 37.76 | -11.01 n.s | 32.09 | 30.24 | -5.76 n.s | 33.85 | 31.41 | -7.21 n.s | 43.10 | 38.61 | -10.42 n.s |
| GY (g/m<sup>2</sup>) | 693.72 | 553.84 | -20.16 ** | 412.28 | 306.39 | -25.68 * | 412.68 | 384.40 | -6.85 n.s | 537.40 | 326.21 | -39.30 n.s |
| GPC (%) | 12.40 | 13.45 | 8.47 * | 14.37 | 15.31 | 6.54 n.s | 14.42 | 15.93 | 10.47 n.s | 11.91 | 13.51 | 13.43 n.s |
| ZEL (%) | 31.66 | 33.18 | 4.80 n.s | 31.91 | 32.93 | 3.20 n.s | 34.08 | 35.64 | 4.58 * | 33.69 | 32.33 | -4.04 n.s |
| GH (%) | 63.67 | 65.35 | 2.64 n.s | 64.18 | 65.93 | 2.73 n.s | 58.17 | 62.67 | 7.74 * | 60.08 | 64.83 | 7.91 n.s |
| Zn (mg/g) | 0.55 | 0.59 | 7.27 n.s | 0.63 | 0.72 | 14.28 n.s | 0.55 | 0.63 | 14.55 n.s | 0.57 | 0.62 | 8.77 n.s |
| Fe (mg/g) | 0.45 | 0.52 | 15.56 n.s | 0.56 | 0.65 | 16.07 n.s | 0.53 | 0.55 | 3.77 n.s | 0.47 | 0.50 | 6.38 n.s |
| Na<sup>+</sup> (mg/g) | 3.96 | 4.66 | 17.68 n.s | 3.96 | 3.77 | -4.80 n.s | 2.93 | 2.94 | 0.34 n.s | 3.43 | 3.61 | 5.25 n.s |
| K<sup>+</sup> (mg/g) | 6.75 | 7.91 | 17.18 n.s | 7.02 | 7.86 | 11.96 n.s | 6.78 | 7.06 | 4.13 n.s | 7.13 | 7.96 | 11.64 * |
| MOF (%) | 10.33 | 9.84 | -4.74 n.s | 10.69 | 10.16 | -4.96 n.s | 9.93 | 9.29 | -6.45 n.s | 10.33 | 9.86 | -4.55 n.s |
| WAF (%) | 67.85 | 68.50 | 0.96 n.s | 69.24 | 69.81 | 0.82 n.s | 66.43 | 67.71 | 1.93 n.s | 66.59 | 68.75 | 3.24 n.s |
| WSRC (%) | 105.55 | 124.32 | 17.78 ** | 90.97 | 99.46 | 9.33 ** | 88.23 | 99.43 | 12.69 ** | 90.45 | 114.05 | 26.09 ** |
| SuSRC (%) | 137.24 | 160.32 | 16.82 ** | 124.92 | 135.94 | 8.82 ** | 118.29 | 129.30 | 9.31 ** | 121.31 | 148.54 | 22.45 ** |
| LASRC (%) | 154.31 | 147.30 | -4.54 n.s | 137.51 | 123.44 | -10.23 ** | 120.78 | 105.32 | -12.80 ** | 125.77 | 132.80 | 5.59 * |
| SCSRC (%) | 123.12 | 129.89 | 5.50 n.s | 121.08 | 119.92 | -0.96 n.s | 110.17 | 114.87 | 4.27 n.s | 110.98 | 130.42 | 17.52 ** |

**Table 5-** Continued
| Traits | *T. turgidum ssp. polonicum* |  |  | *T. turgidum ssp. persicum* |  |  | *T. aestivum* |  |  |
| --- | --- | --- | --- | --- | --- | --- | --- | --- | --- |
|  | WW | WS | Change (%) | WW | WS | Change (%) | WW | WS | Change (%) |
| DM (day) | 212.15 | 203.30 | -4.17 * | 212.92 | 207.08 | -2.74 * | 211.92 | 205.50 | -3.03 <sup>n.s</sup> |
| TGW (g) | 43.46 | 39.81 | -8.40 <sup>n.s</sup> | 32.89 | 29.05 | -11.67 * | 40.39 | 34.22 | -15.28 <sup>n.s</sup> |
| GY (g/m<sup>2</sup>) | 434.21 | 286.60 | -33.99 ** | 467.39 | 304.61 | -34.83 * | 685.07 | 576.34 | -15.87 <sup>n.s</sup> |
| GPC (%) | 13.52 | 14.52 | 7.40 * | 14.11 | 15.39 | 9.07 <sup>n.s</sup> | 11.77 | 12.48 | 6.03 <sup>n.s</sup> |
| ZEL (%) | 33.60 | 33.49 | -0.33 <sup>n.s</sup> | 33.28 | 34.75 | 4.42 <sup>n.s</sup> | 34.06 | 32.68 | -4.05 <sup>n.s</sup> |
| GH (%) | 64.15 | 66.75 | 4.05 <sup>n.s</sup> | 62.67 | 64.75 | 3.32 <sup>n.s</sup> | 52.00 | 53.08 | 2.08 <sup>n.s</sup> |
| Zn (mg/g) | 0.64 | 0.69 | 7.81 <sup>n.s</sup> | 0.64 | 0.65 | 1.56 <sup>n.s</sup> | 0.59 | 0.54 | -8.47 <sup>n.s</sup> |
| Fe (mg/g) | 0.55 | 0.59 | 7.27 <sup>n.s</sup> | 0.56 | 0.61 | 8.93 <sup>n.s</sup> | 0.47 | 0.51 | 8.51 <sup>n.s</sup> |
| Na<sup>+</sup> (mg/g) | 3.51 | 3.60 | 2.56 <sup>n.s</sup> | 4.12 | 3.56 | -13.59 <sup>n.s</sup> | 3.10 | 3.05 | -1.61 <sup>n.s</sup> |
| K<sup>+</sup> (mg/g) | 7.07 | 8.00 | 13.15 <sup>n.s</sup> | 6.41 | 7.36 | 14.82 <sup>n.s</sup> | 6.37 | 6.94 | 8.95 <sup>n.s</sup> |
| MOF (%) | 10.22 | 9.81 | -4.01 <sup>n.s</sup> | 10.38 | 9.75 | -6.07 * | 10.23 | 9.79 | -4.30 <sup>n.s</sup> |
| WAF (%) | 69.60 | 69.80 | 0.29 <sup>n.s</sup> | 68.46 | 69.26 | 1.17 <sup>n.s</sup> | 64.34 | 64.91 | 0.88 <sup>n.s</sup> |
| WSRC (%) | 101.64 | 113.88 | 12.04 ** | 104.88 | 115.23 | 9.87 ** | 91.94 | 114.41 | 24.44 ** |
| SuSRC (%) | 136.05 | 152.07 | 11.77 ** | 137.62 | 145.68 | 5.86 * | 121.37 | 147.95 | 21.90 ** |
| LASRC (%) | 145.54 | 140.06 | -3.76 <sup>n.s</sup> | 143.41 | 133.79 | -6.71 * | 122.70 | 116.93 | -4.70 <sup>n.s</sup> |
| SCSRC (%) | 119.33 | 130.95 | 9.74 ** | 133.92 | 140.81 | 5.14 * | 113.21 | 121.76 | 7.55 * |

\* and \*\* show significance at the 0.05 and 0.01 probability levels, respectively.
n.s: not significant
DM, days to maturity; Fe, Iron content; GH, grain hardness; GPC, grain protein content; GY, grain yield; K<sup>+</sup>, potassium content; LASRC, lactic acid SRC; MOF, moisture of flour; Na<sup>+</sup>, sodium content; SCSRC, sodium carbonate SRC; SuSRC, sucrose SRC; TGW, thousand grain weight; WAF, water absorption of flour; WSRC, water SRC; ZEL, Zeleny index; Zn, zinc content.

Considerable variations in terms of SRC values were found among all studied genotypes. In 2019, WSRC ranged from 82.30 to 118.68% under WW condition and 90.74 to 134.48% under WS. In 2020, it ranged from 83.88 to 146.02% under WW and 96.54 to 181.00% under WS condition. In 2019, SuSRC varied from 112.83 to 147.40% under WW condition and 119.05 to 171.95% under WS condition; while in 2020 its range was from 117.12 to 186.82% under WW condition and 126.96 to 230.23% under WS (Table S1). The range of LASRC was from 116.55 to 169.19% under WW condition, and 76.29 to 150.52% under WS condition, in 2019 and from 112.20 to 203.05% under WW condition, and 88.05 to 193.06% under WS in 2020. In 2019, under WW condition the SCSRC varied from 101.74 to 148.06%, and under WS condition it ranged from 109.09 to 165.49%. In 2020, it varied from 100.36 to 161.86% under WW condition, and 101.22 to 187.17% under WS (Table S1).

Phenotypic and genotypic coefficients of variation (PCV and GCV) for WW and WS conditions are given in Table 4. PCV had a range of 2.04% for DM and 17.59% for TGW under WW condition and 1.87% for DM and 22.87% for Na^+^ under WS. GCV ranged in 1.49% for MOF and 15.76% for TGW under WW condition and 1.47% for DM and 17.05% for Na^+^ under WS treatment. For some traits such as DM, TGW, GPC, ZEL, and WAF, the variations under WW condition were higher than the ones for WS condition (Table 4). For the remaining traits of GY, Na^+^, K^+^, MOF, WSRC, SuSRC, LASRC, and SCSRC, both PCV and GCV were higher under WS condition. Based on GCV, the highest range of genetic variation was observed for GY, and relatively lower ones were detected for DM, GH, Fe, K^+^, and WAF, respectively (Table 4).

Broad-sense heritability estimates and variance components of all measured traits based on all subspecies are given in Table 6. Heritability estimates ranged from 33.78% for MOF and 93.12% for GH, based on the combined data over two years and two moisture environments. Moreover, heritability estimates were also calculated for each moisture environment separately. Under WW condition, this parameter ranged from 34.95% for MOF and 92.51% for GPC; while under WS condition it was from 29.11% for MOF and 83.89% for WAF. Results also revealed that, except for GH, Fe, and LASRC, heritability estimates of all traits were higher under WW environment than the WS one (Table 6).

**Table 6.** Estimates of variance components and broad- sense heritability of the evaluated traits in 36 wheat genotypes under two moisture environments (well-watering and water stress) during 2019 and 2020.

| **Combined** |  | **Traits** |  |  |  |  |  |  |
| --- | --- | --- | --- | --- | --- | --- | --- | --- |
| Variance components | DM | TGW | GY | GPC | ZEL | GH | Zn | Fe |
| $\sigma_g^2$ | 1.74 | 31.42 | 3230.67 | 1.2794 | 0.76 | 28.37 | 0.0021 | 0.0024 |
| $\sigma_{ge}^2$ | 5.46 | -1.78 | 3752.02 | 0.0020 | -0.41 | -5.13 | 0.0004 | 0.0001 |
| $\sigma_{gey}^2$ | -1.14 | 2.00 | 471.48 | 0.2648 | 2.34 | 11.54 | 0.0008 | 0.0002 |
| $\sigma_e^2$ | 7.68 | 17.90 | 12462.11 | 0.1834 | 5.90 | 17.86 | 0.0040 | 0.0084 |
| $\sigma_p^2$ | 33.34 | 38.17 | 6712.39 | 1.3945 | 1.86 | 30.47 | 0.0034 | 0.0039 |
| $h_b^2$ (%) | 80.22 | 82.31 | 48.13 | 91.75 | 41.25 | 93.12 | 60.86 | 60.33 |
| **Well-watering condition** |  |  |  |  |  |  |  |  |
| Variance components | DM | TGW | GY | GPC | ZEL | GH | Zn | Fe |
| $\sigma_g^2$ | 15.28 | 37.66 | 1461.36 | 1.34 | 3.85 | 21.91 | 0.0032 | 0.0024 |
| $\sigma_{gy}^2$ | 3.08 | 9.44 | -5106.89 | 0.12 | 3.16 | 12.10 | 0.0005 | 0.0028 |
| $\sigma_e^2$ | 7.76 | 18.24 | 14065.41 | 0.18 | 7.52 | 28.90 | 0.0026 | 0.0073 |
| $\sigma_p^2$ | 18.76 | 46.94 | 2424.26 | 1.44 | 7.32 | 35.18 | 0.0041 | 0.0056 |
| $h_b^2$ (%) | 81.43 | 80.24 | 60.28 | 92.51 | 52.67 | 62.27 | 78.33 | 42.71 |
| **Water stress condition** |  |  |  |  |  |  |  |  |
| Variance components | DM | TGW | GY | GPC | ZEL | GH | Zn | Fe |
| $\sigma_g^2$ | 9.12 | 21.62 | 3504.04 | 1.23 | 1.61 | 24.59 | 0.0023 | 0.0029 |
| $\sigma_{gy}^2$ | 7.41 | 14.17 | 770.25 | 0.51 | 1.40 | 9.15 | 0.0030 | -0.0005 |
| $\sigma_e^2$ | 7.60 | 17.56 | 10858.80 | 0.18 | 4.29 | 6.81 | 0.0052 | 0.0095 |
| $\sigma_p^2$ | 14.73 | 33.09 | 6603.86 | 1.53 | 3.38 | 30.87 | 0.0051 | 0.0050 |
| $h_b^2$ (%) | 61.96 | 65.32 | 53.06 | 80.44 | 47.56 | 79.66 | 44.81 | 57.80 |

**Table 6– Continued**
| **Combined** |  | **Traits** |  |  |  |  |  |  |
| --- | --- | --- | --- | --- | --- | --- | --- | --- |
| Variance components | Na<sup>+</sup> | K<sup>+</sup> | MOF | WAF | WSRC | SuSRC | LASRC | SCSRC |
| $\sigma_g^2$ | 0.26 | 0.12 | 0.02 | 2.89 | 75.28 | 86.58 | 203.79 | 89.54 |
| $\sigma_{ge}^2$ | 0.07 | -0.06 | -0.02 | 0.11 | 21.42 | 15.34 | 14.14 | 30.99 |
| $\sigma_{gey}^2$ | 0.14 | 0.33 | 0.08 | 0.45 | 80.36 | 65.89 | 73.29 | 23.57 |
| $\sigma_e^2$ | 0.14 | 0.50 | 0.07 | 0.50 | 14.42 | 26.24 | 29.79 | 29.21 |
| $\sigma_p^2$ | 0.49 | 0.26 | 0.06 | 3.15 | 119.98 | 136.07 | 277.05 | 124.54 |
| $h_b^2$ (%) | 52.42 | 46.67 | 33.78 | 91.62 | 62.75 | 63.63 | 73.56 | 71.90 |
| **Well-watering condition** |  |  |  |  |  |  |  |  |
| Variance components | Na<sup>+</sup> | K<sup>+</sup> | MOF | WAF | WSRC | SuSRC | LASRC | SCSRC |
| $\sigma_g^2$ | 0.23 | 0.26 | 0.02 | 3.17 | 76.56 | 75.90 | 176.76 | 92.16 |
| $\sigma_{gy}^2$ | 0.31 | 0.13 | 0.05 | 0.54 | 79.85 | 75.42 | 154.95 | 39.01 |
| $\sigma_e^2$ | 0.07 | 0.49 | 0.08 | 0.38 | 9.98 | 18.06 | 28.31 | 25.53 |
| $\sigma_p^2$ | 0.41 | 0.45 | 0.07 | 3.54 | 118.98 | 118.13 | 261.31 | 118.05 |
| $h_b^2$ (%) | 57.06 | 57.39 | 34.95 | 89.60 | 64.34 | 64.25 | 67.64 | 78.07 |
| **Water stress condition** |  |  |  |  |  |  |  |  |
| Variance components | Na<sup>+</sup> | K<sup>+</sup> | MOF | WAF | WSRC | SuSRC | LASRC | SCSRC |
| $\sigma_g^2$ | 0.44 | 0.37 | 0.05 | 2.08 | 113.63 | 127.41 | 254.57 | 152.28 |
| $\sigma_{gy}^2$ | 0.61 | 0.62 | 0.21 | 0.49 | 132.03 | 150.64 | 186.26 | 71.71 |
| $\sigma_e^2$ | 0.19 | 0.51 | 0.06 | 0.62 | 18.66 | 34.05 | 31.20 | 32.73 |
| $\sigma_p^2$ | 0.79 | 0.81 | 0.17 | 2.48 | 184.31 | 211.24 | 355.50 | 196.31 |
| $h_b^2$ (%) | 55.62 | 46.22 | 29.11 | 83.89 | 61.65 | 60.31 | 71.61 | 77.57 |

DM, days to maturity; Fe, Iron content; GH, grain hardness; GPC, grain protein content; GY, grain yield; K<sup>+</sup>, potassium content; LASRC, lactic acid SRC; MOF, moisture of flour; Na<sup>+</sup>, sodium content; SCSRC, sodium carbonate SRC; SuSRC, sucrose SRC; TGW, thousand grain weight; WAF, water absorption of flour; WSRC, water SRC; ZEL, Zeleny index; Zn, zinc content.

Phenotypic correlation coefficients between different traits based on the average of all subspecies and two years showed that grain yield had significant and negative correlations with GPC, Zn, and Fe under both WW and WS conditions. Under both conditions, grain protein content (GPC) had significant and positive associations with GH, Zn, Fe, and WAF (Table 7).

**Table 7.** Correlation coefficients among morphological, agronomic and quality-related traits of 36 wheat genotypes in well-watering (above diagonal) and water stress (below diagonal) conditions during 2019 and 2020.

|  | DM | TGW | GY | GPC | ZEL | GH | Zn | Fe | Na<sup>+</sup> | K<sup>+</sup> | MOF | WAF | WSRC | SuSRC | LASRC | SCSRC |
| --- | --- | --- | --- | --- | --- | --- | --- | --- | --- | --- | --- | --- | --- | --- | --- | --- |
| DM | 1 | -0.24 | -0.51** | 0.32 | 0.12 | -0.01 | 0.07 | 0.27 | -0.23 | -0.04 | -0.17 | 0.01 | -0.29 | -0.38* | -0.29 | -0.32 |
| TGW | -0.51** | 1 | 0.57** | -0.47** | 0.04 | -0.04 | -0.25 | -0.37* | -0.26 | -0.12 | -0.23 | -0.04 | 0.31 | 0.29 | 0.27 | -0.18 |
| GY | -0.40* | 0.42* | 1 | -0.65** | -0.16 | -0.19 | -0.54** | -0.63** | 0.01 | -0.27 | -0.14 | -0.35* | 0.26 | 0.14 | 0.27 | -0.11 |
| GPC | 0.35* | -0.39* | -0.56** | 1 | 0.19 | 0.34* | 0.47** | 0.78** | -0.05 | 0.15 | -0.01 | 0.57** | -0.15 | 0.01 | -0.05 | 0.13 |
| ZEL | 0.38* | -0.05 | 0.04 | 0.49** | 1 | -0.17 | 0.16 | 0.19 | -0.61** | -0.27 | -0.43** | -0.14 | -0.43** | -0.35* | -0.53** | -0.41* |
| GH | -0.16 | 0.23 | -0.04 | 0.44** | 0.16 | 1 | 0.06 | 0.25 | 0.28 | 0.29 | -0.06 | 0.85** | 0.24 | 0.35* | 0.46** | 0.25 |
| Zn | 0.25 | -0.36* | -0.71** | 0.63** | 0.09 | 0.19 | 1 | 0.74** | -0.02 | 0.06 | 0.02 | 0.25 | -0.01 | 0.08 | -0.10 | 0.17 |
| Fe | 0.18 | -0.37* | -0.43** | 0.61** | 0.18 | 0.13 | 0.67** | 1 | -0.08 | 0.17 | -0.08 | 0.45** | -0.12 | 0.03 | -0.08 | 0.10 |
| Na<sup>+</sup> | -0.34* | 0.01 | 0.30 | -0.32 | -0.39* | 0.30 | -0.11 | -0.17 | 1 | 0.49** | 0.37* | 0.29 | 0.63** | 0.60** | 0.70** | 0.78** |
| K<sup>+</sup> | -0.21 | 0.03 | 0.07 | -0.01 | -0.24 | 0.43** | 0.20 | 0.03 | 0.68** | 1 | 0.05 | 0.44** | 0.23 | 0.37* | 0.43** | 0.46** |
| MOF | -0.11 | -0.20 | -0.26 | -0.15 | -0.46** | -0.08 | 0.37* | 0.31 | 0.26 | 0.18 | 1 | 0.12 | -0.04 | 0.06 | 0.09 | 0.26 |
| WAF | -0.09 | 0.10 | -0.24 | 0.55** | 0.16 | 0.94** | 0.37* | 0.29 | 0.21 | 0.42* | 0.08 | 1 | 0.28 | 0.48** | 0.53** | 0.39* |
| WSRC | -0.33* | 0.21 | 0.26 | -0.42* | -0.21 | 0.12 | -0.29 | -0.32 | 0.66** | 0.49** | 0.06 | -0.02 | 1 | 0.85** | 0.87** | 0.71** |
| SuSRC | -0.36* | 0.26 | 0.23 | -0.34* | -0.22 | 0.22 | -0.16 | -0.20 | 0.64** | 0.50** | 0.11 | 0.08 | 0.94** | 1 | 0.89** | 0.79** |
| LASRC | -0.45** | 0.31 | 0.16 | -0.21 | -0.21 | 0.53** | -0.06 | -0.17 | 0.73** | 0.61** | 0.19 | 0.42* | 0.84** | 0.88** | 1 | 0.73** |
| SCSRC | -0.17 | -0.10 | -0.20 | 0.10 | -0.06 | 0.31 | 0.12 | 0.00 | 0.43** | 0.51** | 0.01 | 0.28 | 0.72** | 0.73** | 0.68** | 1 |

\* and \*\* show significance at the 0.05 and 0.01 probability levels, respectively.
DM, days to maturity; Fe, Iron content; GH, grain hardness; GPC, grain protein content; GY, grain yield; K<sup>+</sup>, potassium content; LASRC, lactic acid SRC; MOF, moisture of flour; Na<sup>+</sup>, sodium content; SCSRC, sodium carbonate SRC; SuSRC, sucrose SRC; TGW, thousand grain weight; WAF, water absorption of flour; WSRC, water SRC; ZEL, Zeleny index; Zn, zinc content.

Under WS, GPC and DM showed significant and positive correlations with ZEL and negative correlations with WSRC and SuSRC; while these correlations were not significant under WW condition. Under WW condition, ZEL had significant and negative associations with WSRC, SuSRC, LASRC, and SCSRC; while this was not the case under WS condition. Under both conditions, each of the four SRC-related traits had significant and positive correlations with Na^+^, K^+^, and the three other SRC traits. Under WW condition, SuSRC was positively correlated with GH and WAF, and SCSRC was associated with WAF. Moreover, LASRC was positively associated with GH and WAF under both conditions (Table 7).

A multivariate technique (i.e. PCA) was conducted for WW and WS conditions to acquire a unifying view of the relationship between the SRC and phenological, agronomic and quality-related traits. It was shown that the first two principal components explained 57 and 56% of the total variation incorporated in the data under the WW and WS conditions, respectively (Fig. 1). According to the differences in plant architecture of tetraploid wheat subspecies and also the results of PCA, the measured traits were divided into several groups based on the correlation matrix and cosine of the angles between the vectors, each of which was associated with specific subspecies. Traits related to nutritional quality, PGP, Fe, and Zn had a strong and positive correlation with each other, and the majority of genotypes of emmer wheat subspecies (*T. turgidum* ssp. *dicoccum*) including G9, G10, G11, G12, G25, and G30 had the higher values for these traits under both moisture conditions (Fig 1). In addition under both moisture conditions, the yield-related traits of TGW and GY showed a strong and positive correlation and half of the genotypes of durum wheat subspecies (*T. turgidum* ssp. *durum*), including Iranian genotypes (i.e. G1, G2, G3, and G4) and International Research Centers CIMMYT (i.e. G5), and ICARDA (i.e. G6) had the higher values for these traits. Moreover, traits related to solvent retention capacity, WSRC, SuSRC, LASRC, and SCSRC were positively and strongly correlated with GH and WAF under both moisture conditions. In these two groups of traits, *T. turgidum* ssp. *durum* (genotypes G7, G8, G27, and G36), *T. turgidum* ssp. *polonicum* (genotype G21), and *T. turgidum* ssp. *persicum* (genotype G31) recorded the higher values for these traits (Fig. 1).

**Fig. 1.**
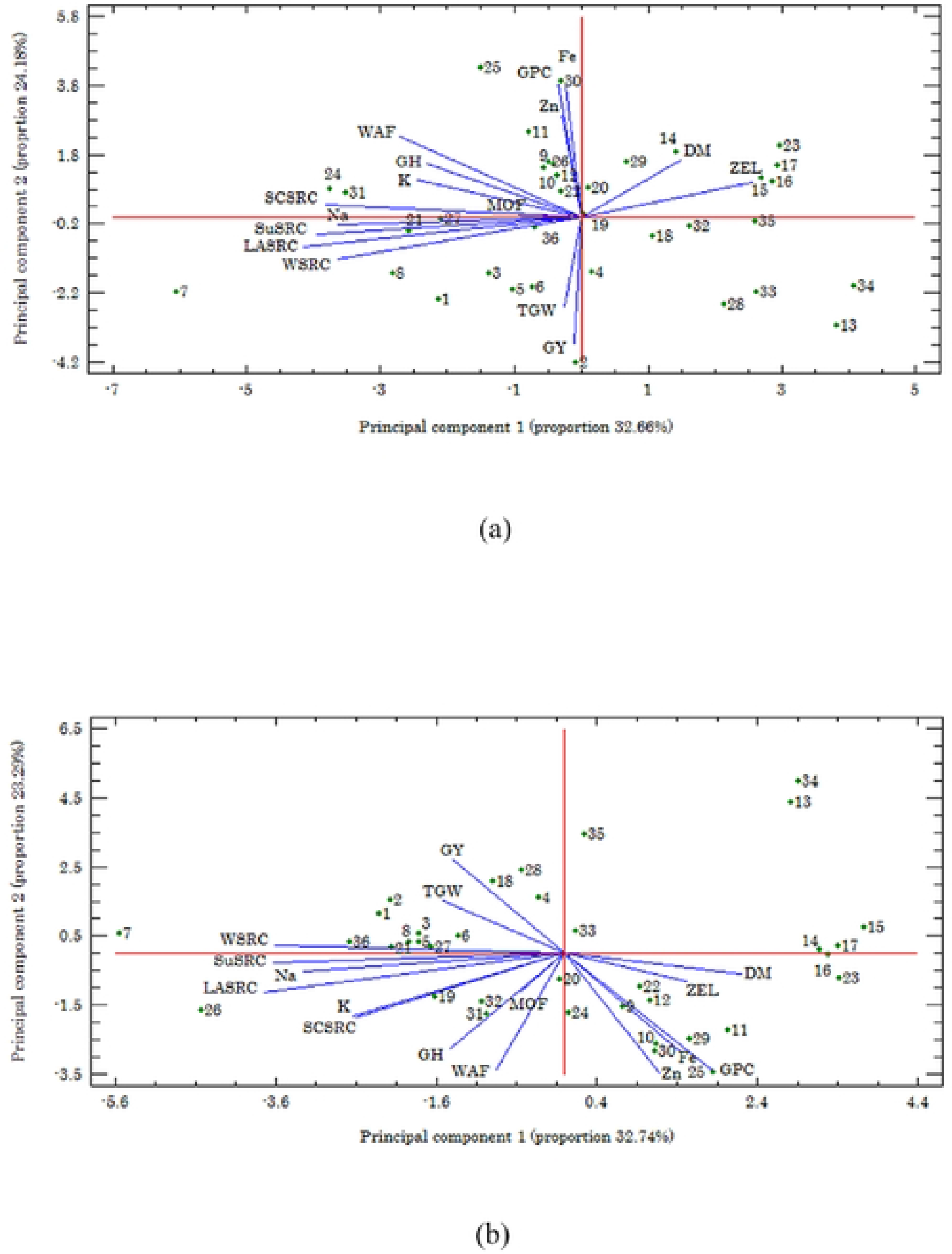
Distribution of the first two principal components (PC) of agronomic, quality, and SRC-related traits in 36 genotypes of wheat belonged to six different subspecies under (a) well-watering, and (b) water stress condition. DM, days to maturity; Fe, Iron content; GH, grain hardiness; GPC, grain protein content; GY, grain yield; K^+^, potassium content; LASRC, lactic acid SRC; MOF, moisture of flour; Na^+^ sodium content; SCSRC, sodium carbonate SRC; SuSRC, sucrose SRC; TGW, thousand grain weight; WAF, water absorption of flour; WSRC, water SRC; ZEL, Zeleny index; Zn, zinc content.

## Discussion

A significant increase in food production is required to meet the food demand of the growing world population. Abiotic stresses are major constraints to crop production and food security worldwide. The situation has aggravated due to the drastic and rapid changes in global climate [13]. Moreover, inter- and intraspecific variability gives cultivated crops the ability to adapt to changes and stabilize production [14]. Landraces are unexploited genetic resources for various agronomic and quality traits contributing tolerance to abiotic stresses. In the present study, considerable genetic variations were observed among wheat genotypes for all of the measured traits, emphasizing the high potential within and among the studied subspecies for improving these traits through targeted selection in breeding programs. Large variations in grain Zn and Fe concentrations among wheat genotypes have also been already reported, suggesting that there is a genetic potential to enhance the levels of these in edible parts to meet human nutritional requirements through biofortification programs [15].

Comparison among the studied subspecies revealed that *T. durum* which had the highest GY, Na^+^, WSRC, SuSRC, and LASRC, and *T. dicoccum* which had the highest GH, Zn, Fe, MOF, and WAF are promising candidates in terms of quality-related traits. Therefore, they can be used in the breeding programs to improve the quality of bread wheat (*T. aestivum*). Moreover, *T. turgidum* ssp. *oriental* with the highest values of GPC and ZEL index can effectively increase the protein content of bread wheat. The high grain quality of emmer is mainly conditioned by high protein content. The high protein content in the emmer grain significantly affects the yield of protein per unit area which can be even higher than that in bread wheat. In the present study, protein content in *dicoccum* wheat ranged from 13.71% to 17.34%. Suchowilska et al. [16] reported high protein content of 22.7% in *T. dicoccum* wheat from Poland.

In addition, among the studied genotypes of all subspecies, the higher values of Zn and Fe were observed for G25, G29, and G30; and the lower values were detected for G7 and G13. Insufficient supply of Zn and Fe is a widespread mineral malnutrition problem among the world’s population, which compromises the immune system and retards development in infancy. Moreover, Fe deficiency can cause nutritional anemia, problem pregnancies, stunted growth, lower resistance to infections, long-term impairment in mental function, decreased productivity, and impaired neural development. While, growth retardation, delayed skeletal and sexual maturity, dermatitis, alopecia, and increase in susceptibility to infection are the complications of Zinc deficiency [17]. A sustainable way to improve Zn and Fe nutrition by increasing the density of these micronutrients in cereal grains is through plant breeding and genetic engineering techniques considering the presence of proper germplasm.

Moreover, the higher values of LASRC were observed for G7, G8, and G27. The LASRC values are explicitly related to the glutenin content of the flour sample [10]. Hence, the genotypes with higher LASRC values specifically indicated higher glutenin content. The G7, G24, and G31 showed higher SCSRC values than others. The SCSRC values are related to the damaged starch content of the flour [18]. Higher SCSRC values indicated a higher amount of damaged starch produced if moisture in grain decreases, making the grain harder. Hard wheat always produces more damaged starch because more force and pressure are required for grinding damaging the starch granule [19]. The higher values of SuSRC were detected in G7, G24, and G27 which is probably due to the high pentosan content; because the sucrose SRC test increases with pentosan available mainly in the cell walls of plant material [20]. Finally, G5, G7, and G24 showed higher values of WSRC. Water is used as a control solvent, and therefore WSRC is not specifically connected to a certain polymer. In fact, WSRC values describe the overall water holding capacity and are lower compared to the other SRC values [18]. These results indicated that the SRC values had a strong diagnostic potential for wheat genotypes and subspecies quality due to different distribution of grains constituents in different genotypes and subspecies, and can be used to differentiate between them.

Water deficit limits agricultural production by preventing crop plants from expressing their full genetic potential. In the current study, WS significantly influenced most of the evaluated traits. As expected, it caused a significant decline in DM, TGW, GY, and LASRC; and significantly increased GPC, ZEL, GH, WAF, WSRC, SuSRC, and SCSRC in both experimental years compared with WW condition. The study region (Najafabad, Isfahan, Iran) is a hot and dry area where summer temperature reaches as high as 45°C and precipitation is null; therefore, as mentioned above reductions are to be expected in some traits. Moreover, WS reduced grain yield and contributed significantly to the low productivity of wheat. This may be ascribed to the stomatal closure in response to low water potential of the soil, decreased rate of photosynthesis, disturbed assimilate partitioning, and disturbance in the grain filling period [13].

In this study, wide genetic variation was observed for all traits within and between subspecies, indicating a high potential for genetic gain through selection in the studied germplasm. The smaller difference between PCV and GCV will lead to more gain through selection, because it shows the more negligible effect of the environment and as a result higher heritability. In the present study, the smaller differences between these two coefficients were observed for DM, GPC, MOF, WAF, and SCSRC, indicating that more gain will result through selecting for these traits.

Estimation of heritability, a measure of the phenotypic variance attributable to genetic variance available to the plant breeders for selection, is necessary to design and implement an effective breeding program to maximize genetic improvement. In this study, considering all subspecies, high heritability estimates were obtained for most studied traits (except for GY, ZEL, K^+^, and MOF) under both WW and WS conditions, suggesting the presence of some major genes or QTLs affecting them. Traits with high heritability could be improved by recurrent or mass selection [5]. These results were consistent with those reported elsewhere in wheat [e.g., 21], which identified some of the major QTLs encoding for functional genes controlling most agronomic traits under WS. However, no literature was found on the heritability of quality-related traits, especially SRC traits in wheat. For most of the traits having high heritability estimates, three subspecies of *T. durum*, *T. polonicum,* and *T. persicum*, showed high genetic variance; therefore, it seems that these subspecies also have high heritabilities for the related traits. Moreover, for GH and WAF with the heritability estimates of 93.12 and 91.62% respectively, two subspecies of *T. durum* and *T. turgidum*, and also *T. aestivum* had high genetic variance and therefore had probably high heritability estimates for these traits, too.

For most of the measured traits, broad-sense heritability was higher in WW condition than those in WS, which were advantageous for successful selection in achieving genetic progress and indicates that phenotypic selection under WW condition would be more effective than WS condition. As different genes may contribute to the same trait in different environments therefore, changes in heritability would seem likely to occur with increased or decreased stress [22]. Low heritability estimates were obtained for the most economically important trait of GY, which resulted in a lower chance for improving this trait through phenotypic selection. As grain yield is a complex trait controlled by many genes, breeders often use indirect selection and well-correlated attributes with the yield to improve it [23]. Several studies have attempted to estimate the heritability of important economic traits that directly affect yield response in wheat, particularly under water-stressed and non-stressed conditions [e.g., 24]. In the present study, most of the evaluated traits had higher heritability estimates than grain yield. Therefore, determining the relationship between grain yield and these traits could lead to effective criteria for indirect selection.

Correlation analysis linked to essential traits is a valuable and conclusive analysis for identifying selection criteria for improving yield potential and developing better cultivars. Quality-related traits, with easy measurements correlated with complex traits such as grain yield, could also make genotype selection more impressive [25]. The phenotypic correlation coefficients based on the average of all subspecies showed significant and negative associations between days to maturity (DM) and grain yield, indicating that breeding for high-yielding and early-maturing plants can be achieved by manipulating wheat phenology. Breeding novel wheat genotypes with early flowering and maturity is an important objective in wheat breeding programs [26]. The focus should be on the developing early-maturing wheat genotypes as an adaptive mechanism for environments experiencing terminal heat and water stress [27]. However, such genotypes should have faster growth rates and accumulate sufficient biomass production in shorter times to increase grain yield potential. Grain yield (GY) was negatively and significantly associated with GPC, Zn, and Fe under WW and WS conditions. These relationships were verified by PCA method and indicated that any increase in these traits might be associated with the decrease of grain yield and vice versa. The negative correlations of DM and ZEL with SRC-related traits (WSRC, SuSRC, LASRC, and SCSRC), GH, Na^+^, K^+^, MOF, and WAF under both water conditions suggested that selection for earliness and low ZEL index can indirectly improve the quality of grain wheat. The significant and positive correlations of GPC with GH, WAF, Zn, and Fe showed that the more grain protein results in more grain quality and hardiness. Grain hardness is a major determinant for what products that wheat can make, with hard grain being primarily used for bread-making [28]. On the other hand, the negative correlation of DM with WSRC and SuSRC showed that the more early-maturing genotypes had higher water absorption and pentosan, and therefore are suitable for bread-making. In contrast, late-maturing genotypes had lower water absorption and pentosan, and therefore are ideal for cookie and cracker production. Generally, soft wheat products such as cookies and crackers require flours with low water absorption. In contrast, hard wheat products such as bread require flours with high water absorption. These factors suggest that pentosan plays a detrimental role in the production of cookies and crackers [29] but a beneficial role in bread-making. Results also showed that WAF had significant and positive associations with GH, SuSRC, and SCSRC, indicating that any increase in the grain hardiness, pentosan, and damaged starch may be associated with an increase in the water absorption of flour. Traits of WSRC, SCSRC, LASRC, and SuSRC were grouped close in PCA, indicating a positive correlation between these four SRC variables. These results were in line with earlier studies that also observed strong correlations between these four SRC values [30]. All these variables indicated the flour water holding capacity.

## Conclusion

In conclusion, the substantial genetic variation observed for all measured traits points to the ample resources of untapped genetic variation among the landraces of wheat, which can be used to improve the stated traits in wheat germplasm through targeted selection in future breeding programs. Results showed that WS could greatly influence agronomic and quality-related traits and thus affect wheat grain yield and flour quality. According to the results, WS had adverse effects on DM, TGW, GY, and LASRC; while significantly increased GPC, ZEL, GH, WAF, WSRC, SuSRC, and SCSRC compared to the control condition. Moreover, because of the moderately low broad-sense heritability for grain yield, both genetic and non-genetic effects played a role in controlling this trait. Therefore, selection based on an index, which is a weighted linear combination of several traits, may be more effective to improve grain yield in recurrent selection programs. The correlation coefficients revealed that the early-maturing genotypes had higher water absorption and pentosan, and therefore are suitable for bread-making. In contrast, late-maturing genotypes are ideal for cookie and cracker production. Based on the association of different traits with SRC values and other quality-related traits, preferable genotypes were identified by the biplot method, which are useful to develop genetic populations for breeding studies of grain quality and functional properties of flour in wheat.

## Acknowledgments

We would like to thank the Isfahan University of Technology (IUT) for supporting this work.

## Supporting information captions

**S1 Table. Mean comparisons of different traits evaluated on 36 genotypes of wheat during two years (2019 and 2020) under well-watering (WW) and water stress (WS) conditions.** (DOC)

**S1 Raw data**

(XLS)

